# The Suppression of *Plasmodium berghei* in *Anopheles coluzzii* infected later with *Vavraia culicis*

**DOI:** 10.1101/2023.02.05.527158

**Authors:** Nayna Vyas-Patel

**Affiliations:** Imperial College London

**Keywords:** Microsporidia, *Vavraia culicis*, Mosquito, *Anopheles coluzzii*, *Plasmodium berghei*, larval host age, spore count, double infections

## Abstract

The suppression of malaria parasites in the presence of an existing microsporidian infection of *Vavraia culicis*, was examined using fluorescent *Plasmodium berghei*. Younger hosts infected with *V. culicis* had a greater suppressive effect on the subsequent development of *P. berghei* than older infected hosts. This effect was density dependant on the numbers of microsporidian spores present, consequently it was also dependant on the age at which mosquitoes were infected with microsporidia, as larvae infected later harboured comparatively fewer *V. culicis* spores. The timing of primary and secondary infections of larval and adult mosquitoes (host age) with different parasites affects parasite development and hence disease outcomes. The use of green fluorescent *P. berghei* enabled easier and rapid visualisation and separation of malaria oocysts from *V. culicis* spores. Different degrees of melanisation of *V. culicis* spores was seen in a number of hosts and merits further investigation.

## Introduction

Infection with microsporidia renders the mosquito less hospitable to the development of later *Plasmodium* infections, (Garnham PCC 1956, Bano L 1958, Bray RS 1958, Fox RM & Weiser J 1959, Hulls RH 1971, Savage KE et al 1972, Ward RA & Savage KE 1972, Gajana A et al 1979, Schenker W et al 1992, Margos G et al 1992, Weiser J & ŽIŽKA Z 2004, Herren JK et al 2020). The first recorded observation of this effect came from infections of laboratory colonies where it proved difficult to infect mosquitoes harbouring microsporidia, with malaria parasites (Garnham PCC 1956). It was noted that ‘’In experimental malaria work, where microsporidian-infected mosquitoes are employed as the vector, there may be interference with the normal development of the oocyst and sporozoites’’ (Garnham PCC 1956). This was followed by numerous reports where infections of larval mosquitoes with microsporidians, then later with malaria parasites, inhibited the development of the second infection i.e., the malaria parasite, both in terms of numbers of oocysts produced and their size. These reports were largely from inadvertent infections of microsporidia in laboratory reared mosquito colonies. Infections of two day old mosquito larvae with the microsporidian *Vavraia culicis* had a suppressive effect on the development of *Plasmodium berghei* (Bargielowski I & Koella JC 2009). The inhibitory effects of one organism on the development of another is not merely restricted to microsporidia, reports examining field collected anophelines indicated that the composition and diversity of the mosquito microbiome could also have a similar effect (Tainchum K et al, 2020; Cirimotich CM et al 2010). Another example is that of filarial nematode infections negatively impacting the development of plasmodium in mosquitoes, leading to the conclusion that treating the filarial nematode could result in the rise of malaria infections (Aliota MT et al 2011).

A study examining the effects of host age on *V. culicis* indicated that infecting older larval mosquitoes led to the production of comparatively fewer spores (Vyas-Patel N, 2020). It was queried (Vyas-Patel N, 2020) whether infecting older larvae could also bring about the same suppressive effect on the development of the malaria parasite, as found by Bargielowski I & Koella JC (2009) and Vyas-Patel N (2020), from early infected larvae. Could the suppressive effect of microsporidia be dependent on the extent of infection i.e., the numbers of microsporidian spores present in a host? The suppressive effect of *V. culicis* on Plasmodium parasites has been shown to be the result of immune priming (Bargielowski I & Koella JC 2009), it could also be a consequence of the numbers of spores present, a physical, nutritional, or metabolic reaction as well as priming of the host’s immune system. Or the result of a combination of factors. It has been reported that the presence of microsporidia could lead to faster moulting and development of infected hosts (Agnew P et al 1990); could microsporidian infection also speed up and affect any subsequent infections with *Plasmodia*, leading to missed observations of oocysts as they quickly developed into the sporozoites stage? It was known that infecting older mosquito larvae with *V. culicis* would present an altered host environment for the development of any subsequent *Plasmodium berghei*, compared to younger larval infections (Vyas-Patel N, 2020). If older and younger larval mosquitoes were infected with *V. culicis*, then subsequently infected with *P. berghei* as adults on the same day, it would give some indication of the suppressive effects of microsporidia when they inhabit larvae at different ages as the numbers of *V. culicis* spores would comparatively be fewer in the older infected larvae (Vyas-Patel N, 2020). Furthermore, the nutritional and immune status of the host would be different in larvae infected at an older age. A study was therefore undertaken to ascertain if older mosquito larval infections with *V. culicis* could still exert a suppressive effect on the development of subsequent infections with *P. berghei*.

## Materials & Method

### Experiments 1 & 2

Mosquito larvae were infected with 20,000 spores per larva, on day 2 for Experiment 1 and day 2 or day 5 for Experiment 2, post hatching, with *V. culicis*, then subsequently infected with *P. berghei* as adults and compared with mosquitoes not infected with microsporidia, but only infected with *P. berghei* – the controls. Infections of adults with *P. berghei* was carried out on five day old female adults. From day 10 until day 23 post infection, mosquito guts and salivary glands were dissected and examined for ookinetes and sporozoites. Photographs were taken of dissected guts and salivary glands. The different experimental groups were kept in separate cages. The aim of Experiment 1 was to determine if prior infection with microsporidia could speed up the development of *P. berghei*, in which case sporozoites would be seen earlier in doubly infected mosquitoes. In the second study, Experiment 2, mosquito larvae were infected with *V. culicis* on day 2 and a separate group of mosquitoes on day 5 post hatching. Subsequently, both groups were infected with *P. berghei* as adults on day 5 post emergence, to determine if infections of older larvae with microsporidia had the same suppressive effect on the development of the malaria parasite, *P. berghei*.

### Mosquito Rearing and Microsporidian (*V. culicis)* culture

*A. coluzzii* mosquitoes originating from the Cameroons were reared as described in Vyas-Patel N, (2020). Cultures of *V. culicis floridensis*, originating from USDA Gainesville, USA, courtesy of JJ Becnel, were maintained as described in Vyas-Patel N, (2020).

### *Plamodium berghei* ookinete culture

*Plasmodium berghei*, a murine parasite first cultured from forest dwelling, African thicket rats (Natarajan R et al, 2002) were employed. The *P. berghei* ookinetes used were from a transgenic cell line, generated to express green fluorescent protein (GFP) throughout the life cycle of the parasite and were produced using non-viral transfection technology, Nucleofactor^R^ as described by Janse CJ et al, (2006). These fluorescent *P. berghei* cultures were maintained in the laboratory of Prof. R Sinden, Imperial College, London (ICL), using the methodology described by Sinden RE (2002).

### *Plasmodium berghei* infections

Ookinetes were transported from Prof. Sinden’s laboratory at Imperial College London (ICL) to the college campus at Silwood Park, Ascot. The ookinete rich blood was centrifuged at 3000 rpm, for 10 minutes and the supernatant removed. The precipitated ookinetes were then quantified using a haemocytometer under a phase contrast microscope. The remaining blood (precipitate containing ookinetes) was mixed with naive mouse blood and calibrated to deliver a concentration of 800 ookinetes/μl. Mosquitoes were introduced to the infected blood by means of membrane feeders specially produced by Glass Precision Engineering Ltd. (GPE Scientific, product code SP 3046). Using a syringe, 300μl of the prepared infected blood was introduced into the top of the feeder. The bottom of the feeder, which was covered with stretched Parafilm was gently placed on top of the mosquito cage containing experimental mosquitoes to be infected, so that the Parafilm only touched the outside of the mosquito netting. Experimental mosquitoes were allowed to feed for an hour, after which the membrane feeders were detached. Mosquitoes that had visibly fed and had distended abdomens were removed and reared in separate cages for each of the groups. They were allowed access to a sugar solution and dissected and examined for oocysts daily from day 10 to 15 (Experiment 1, to determine if there was a speeding up of oocyst development) and day 10 to 23 (Experiment 2, to determine if age at infection with microsporidia affected oocyst development).

### Experimental Design

One thousand five hundred, newly hatched mosquito larvae were individually placed in the well of a 12 well plate; one larva per well, in 2 ml of deionised water per well. Falcon Multiwell™ 12 well plates, (Becton Dickinson) were used. Row one of each plate was infected on day 2 post hatching with 20,000 spores of *V. culicis* as described in Vyas-Patel N (2020). Row two of each plate was similarly infected, but on day 5 post hatching. The remaining row was left uninfected and was the larval control. The treatment rows were randomly allocated in subsequent plates. Mosquito larvae were fed with an increasing amount of food daily, as described in Vyas-Patel N (2020). Larvae were reared in a sealed insectary, kept at a temperature of 26°C (+/−1), with a 12:12 hour light: dark cycle and a humidity of 70% (+/−5). On pupation, each pupa was placed individually in 3 ml of deionised water into a netting covered 50ml Falcon tube and allowed to emerge into adults.

On emergence (usually the next day), each adult mosquito was sexed and males and females were placed separately into labelled cages, around 60 to 80 mosquitoes per cage and supplied with an unlimited amount of sucrose solution. On day 4 post emergence, the female adults were transferred to another rearing room maintained at 19°C, 12:12 hour light: dark cycle, relative humidity of 70% and were deprived of sucrose for 24 hours to encourage feeding the next day (following the advice from ICL, S. Kensington). *P. berghei* maturation is triggered by the drop in temperature from the endothermic mouse body to the ambient temperature and thicket rodents being forest dwelling species, *P. berghei* develops best at the lower 19°C temperature. On day 5 post emergence, adult female mosquitoes were infected with *P. berghei* ookinetes via a membrane blood feeder. Mosquitoes were examined to ensure that their abdomens were full or half full of blood.

Unfed mosquitoes were removed from the cages and fed mosquitoes were allowed access to unlimited amounts of sucrose solution and extra humidity in the form of water-soaked filter paper placed inside the cage. Mosquitoes were dissected and examined for the presence of oocysts and sporozoites each day from day 10 until day 15 (Experiment 1) and day 23 (Experiment 2) post infection.

### Mosquito Dissections

Mosquitoes to be dissected were transferred into cups and placed on ice to anaesthetise the adults. The salivary glands and guts were dissected as described by Service MV (1980) and examined initially under a phase contrast light microscope to determine that doubly infected mosquitoes did indeed show the presence of *V. culicis* spores from every doubly infected mosquito dissected. Also, to determine the absence of *V. culicis* from control mosquitoes, to rule out any kind of observational error. This was followed by examination under the fluorescence microscope to determine the presence/absence of oocysts from the gut and to count the number of oocysts from each gut, if present. To discount the possibility that co-infections speeded up *P. berghei* development and hence the production of sporozoites, the salivary glands were also examined under the fluorescence microscope for the presence of any sporozoites from day 10 onwards.

## Results

### Experiment 1: Does infection with microsporidia speed up the development of malaria parasites?

Figure 1 indicates the presence or absence of oocysts from singly and doubly infected adults. Sporozoites were not seen.

**Figure 1:**
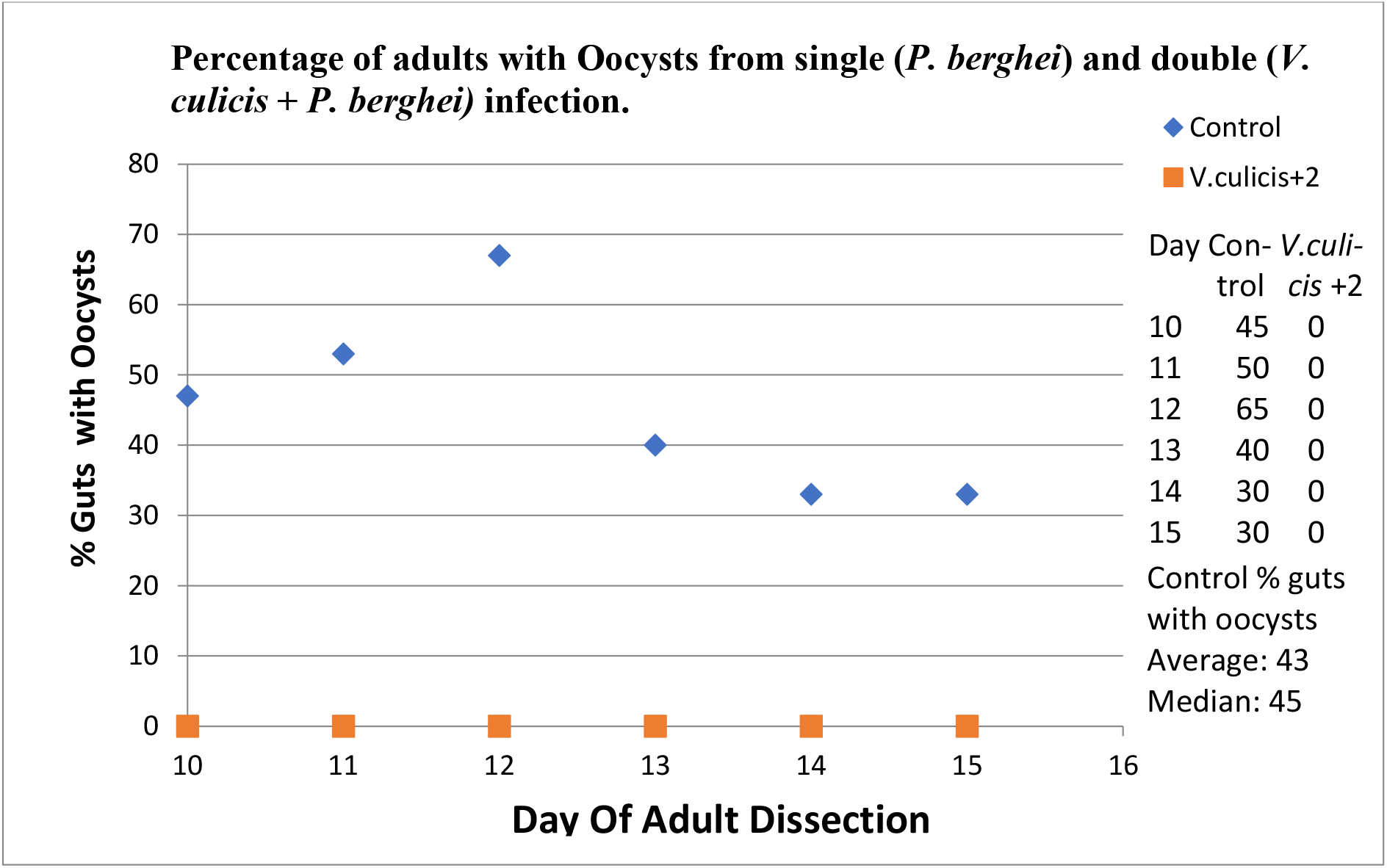
The development of *P. berghei* in the presence and absence (control) of *V. culicis*, day 2 infected larvae.

Control = Adults not infected with *V. culicis*; but singly infected with *P. berghei*.

*V. culicis* + 2 = Mosquitoes infected as larvae, day 2 post hatching, with *V. culicis* and subsequently infected as adults with *P. berghei*.

Figure 1 indicates the complete absence of oocysts from doubly infected (with *V. culicis*, day 2 larvae infections *+ P. berghei*) mosquitoes (0 values) and their presence from singly infected (with *P. berghei*) - the control mosquitoes.

The highest levels of *P. berghei* from single infections (controls) were recorded on day 12; this is within the normal range for peak oocyst formation in single infections with *P. berghei*. The average percentage of mosquitoes infected with oocysts in controls over the 6 days was 43%, the median numbers of mosquitoes infected was also 43% compared to double infections, where it was zero. Counting the numbers of oocysts per host observed, it was noted that the minimum oocyst number per gut was one; the maximum was 27 per gut; the median value was 6 per gut. A host was counted as being positive for oocysts regardless of the numbers of oocysts observed in a gut.

### Experiment 2: Do later infections (of older larvae) with Microsporidia also suppress the development of Malaria parasites?

The second experiment compared the development of oocysts in double infections where the microsporidian infection was introduced early on (day 2 post hatching, younger hosts) and later on (day 6 post hatching, older hosts), with singly infected (only with *P. berghei*) controls. All three of the mosquito groups were infected with *P. berghei* on day 5 post adult emergence. The aim was to determine if the timing/delay of the initial microsporidian infection and hence host age at infection could have any impact on oocyst production in doubly infected mosquitoes. Figure 2 below is a graphical representation of the results, indicating the presence/absence of oocysts from all three treatments.

**Figure 2.**
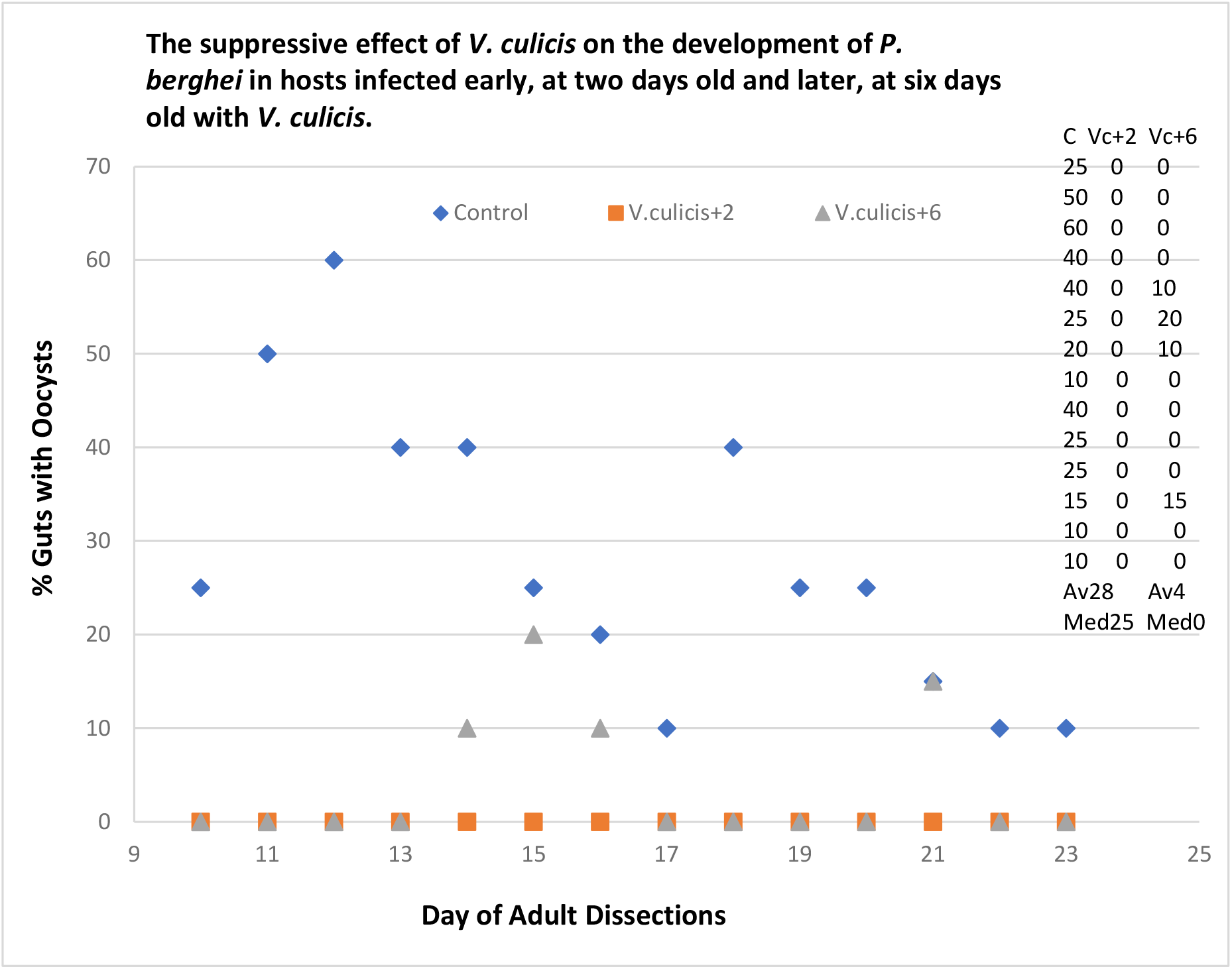
The development of *P. berghei* in younger, day 2 infected larvae (Vc+2), compared with older larvae infected with *V. culicis* on day 6 post hatching (Vc+6); C = Controls (infected only with *P. berghei)*.

Control (C) = Mosquitoes not infected with *V. culicis*; but singly infected with *P. berghei*. *Vc* +2 & Vc+6 = Mosquitoes infected as larvae on day 2 and day 6 post hatching, with *V. culicis* and then infected as adults on day 2 post emergence, with *P. berghei*.

The average percentage of hosts found positive for oocysts in controls was 28%. The median numbers of mosquitoes infected was 25. The average percentage of mosquitoes with oocysts in double infections, day 2 + *V. culicis* was 0, indicating as before, the complete suppression of oocyst development in day 2 infected mosquitoes. The average percentage of mosquitoes with oocysts in double infections, day 6 +*V. culicis* was 4%, median value 0, indicating incomplete suppression of oocyst development from later infected hosts.

Figures 3 & 4 illustrates that under the dissection and light microscopes, a mosquito gut infected with microsporidia can look remarkably similar to one infected by *Plasmodium*. The developing sporophorous vesicles of *V. culicis* look similar to groups of oocysts when they protrude outwards from the gut. Microsporidian species vary in size from 1 to 4 μm; *Plasmodium* oocysts are generally 30 to 40 μm on day 9 post infection. Spores were seen to break out of the gut wall and float into the body cavity. Different degrees of *V. culicis* spore melanisation was seen.

**Figure 3.**
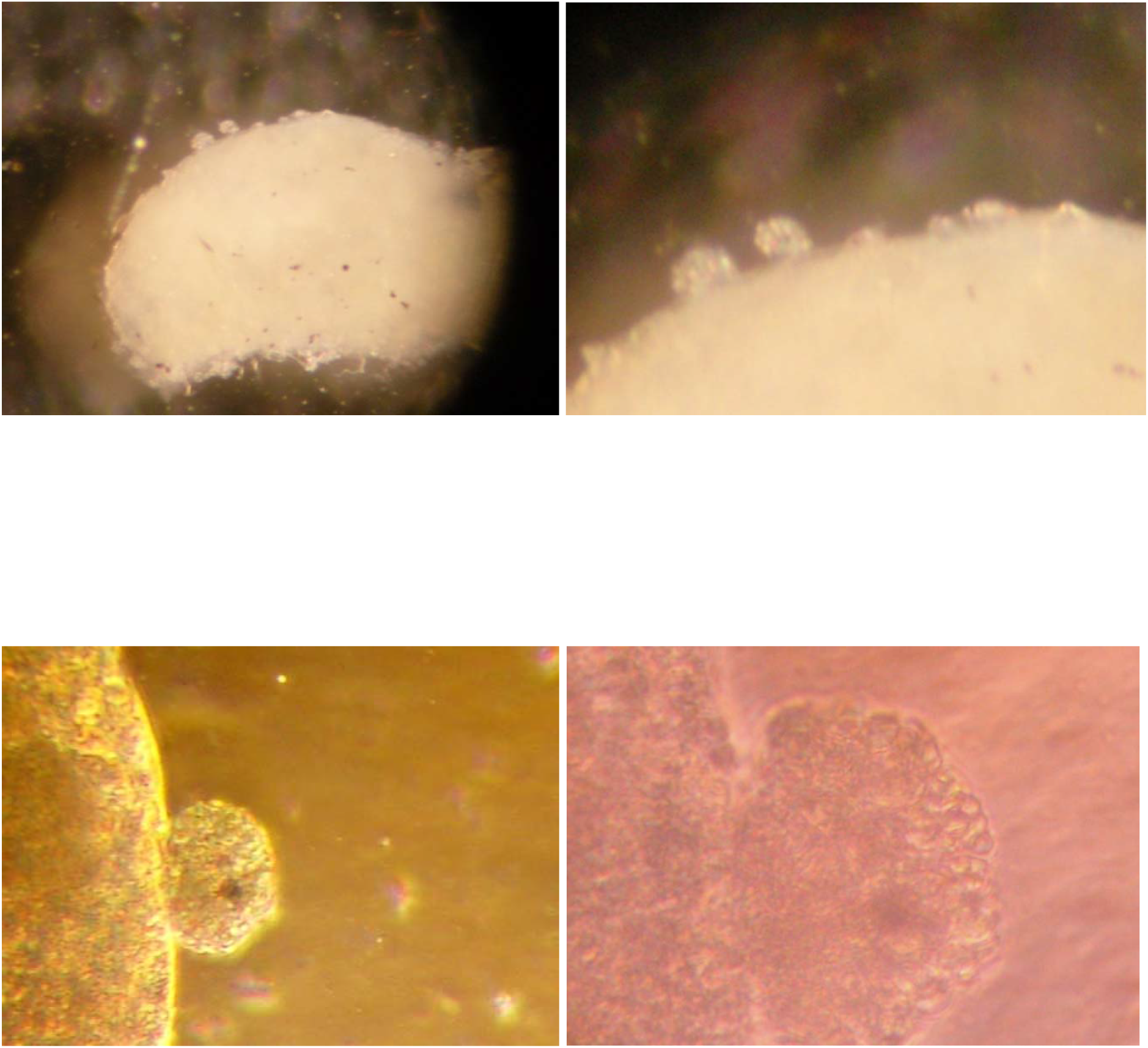
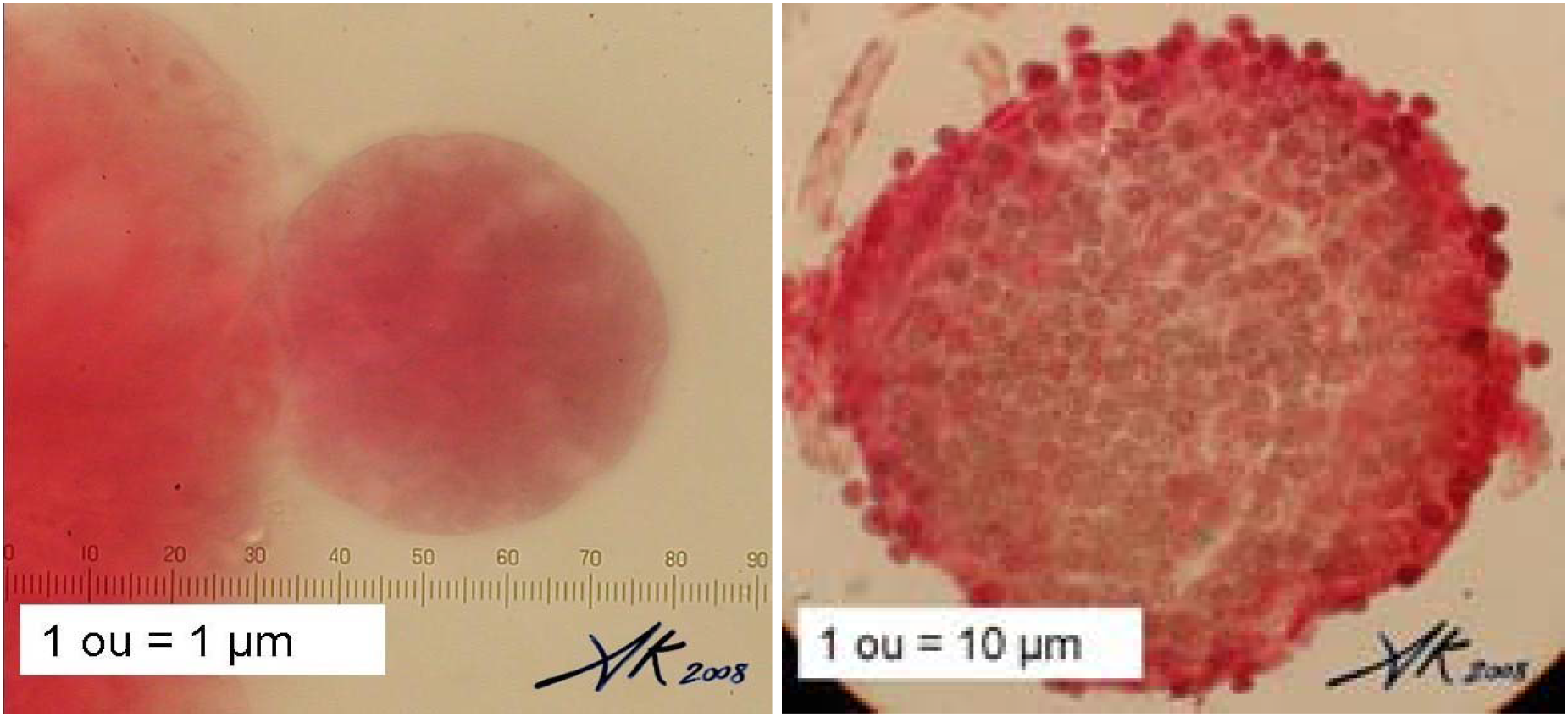
Images of singly (either with *V. culicis* or *P. berghei*) infected mosquito guts using the dissection and phase contrast microscopes. 3a. A mosquito gut heavily & singly infected with *V. culicis*, note the sporophorous vesicles on the outside of the gut. 3b. Close up of sporophorous vesicles on the outside of the mosquito gut. 3c. A sporophorous vesicle, outside of the mosquito gut, under the phase contrast microscope, X200. 3d. The close connection of the protruding sporophorous vesicle with the mosquito gut wall, X400. 3e. *Plasmodium* oocyst on the outside of a mosquito gut, courtesy of Alain Knipes, note the similarity with *V. culicis* sporophorous vesicles. The outline of *Plasmodium* is better defined than *V. culicis* sporophorous vesicles. 3f. Stained *Plasmodium* oocysts on the outside of a mosquito gut, looking remarkably similar to the sporophorous vesicles of *V. culicis*, picture 3a & 3b, above. Photo courtesy of Alain Knipes.

**Figure 4.**
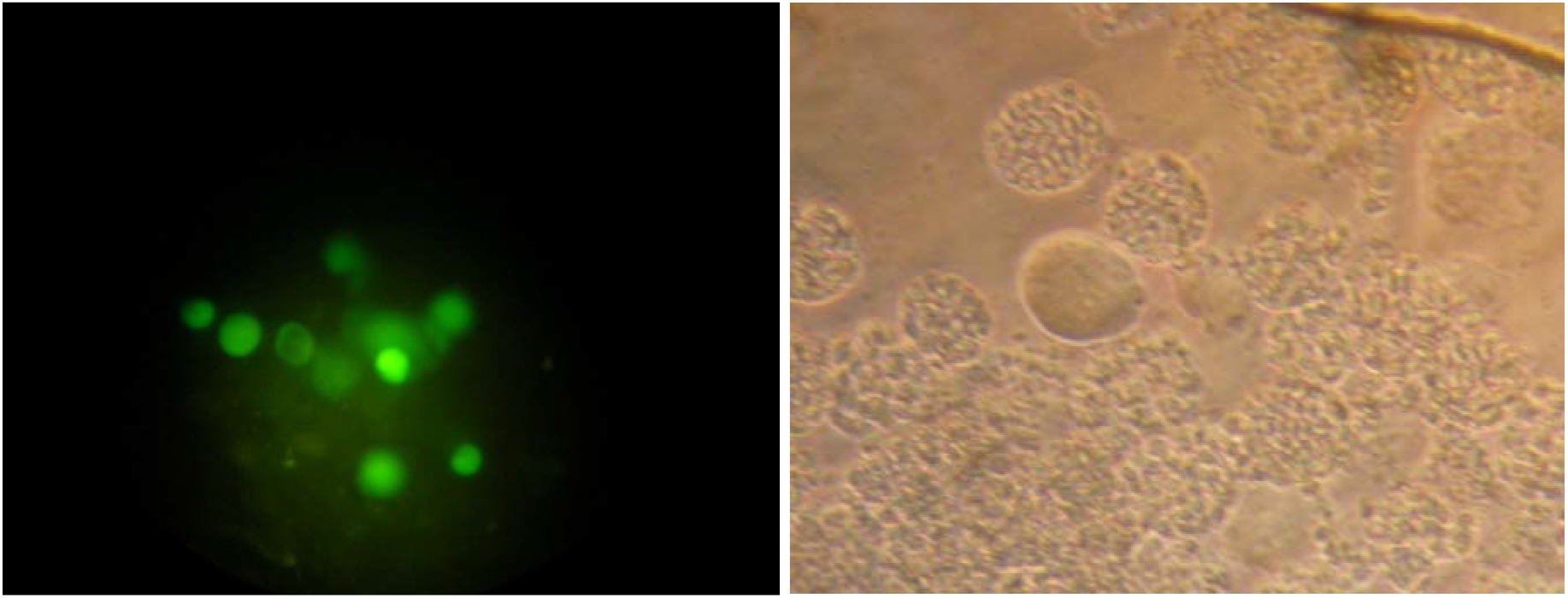
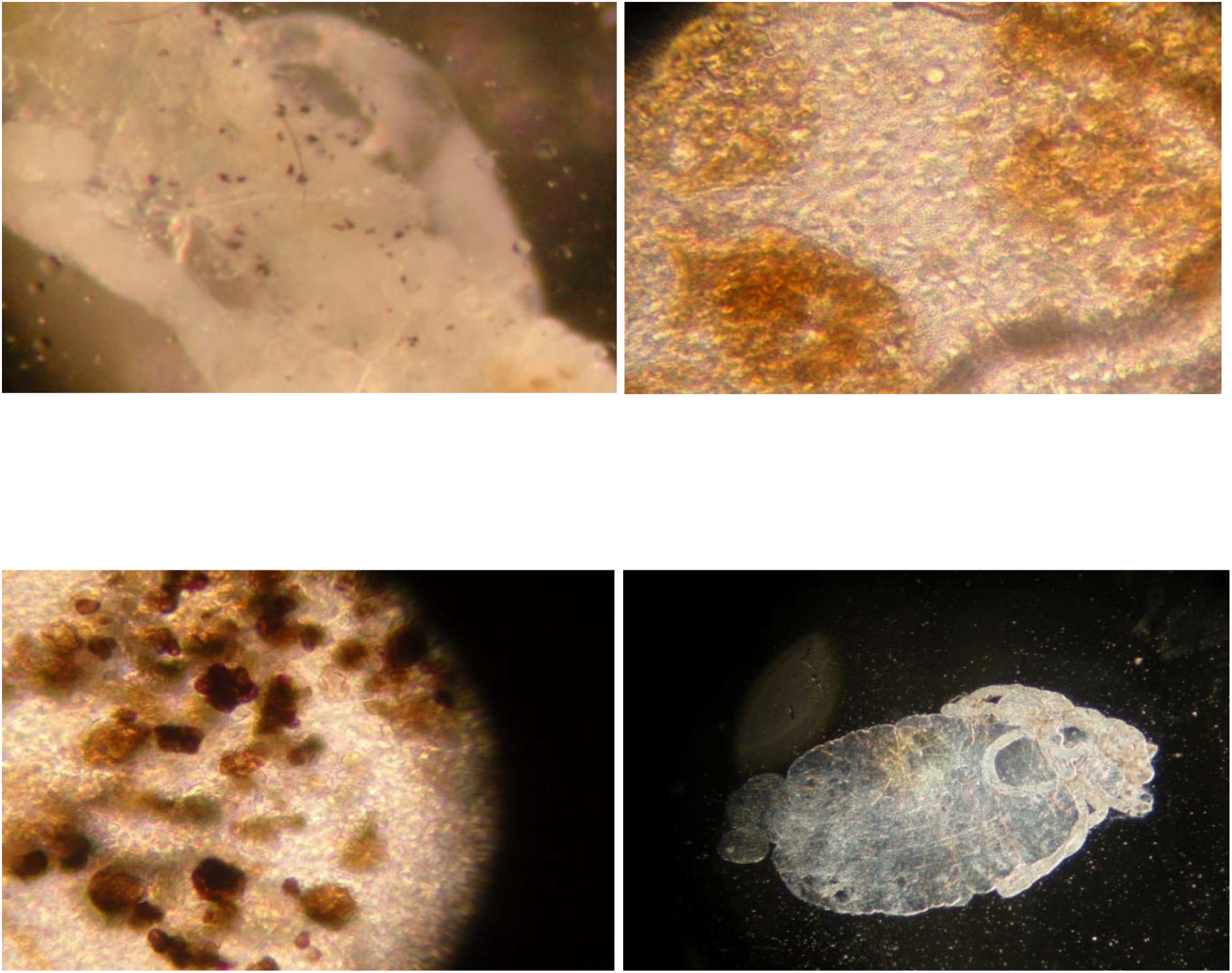
Images of doubly infected mosquitoes (*V. culicis + P. berghei)* using fluorescence and phase contrast microscopes. 4a. *Plasmodium berghei* oocysts, fluorescing; hence easily seen and counted, using the fluorescence microscope. 4b. A doubly infected mosquito gut, phase contrast microscope, X100. Smooth *P. berghei* oocyst in the middle, surrounded by *V. culicis*. 4c. Speckles of melanised spores seen in the mosquito gut under the dissection microscope. 4d. Patches of light melanisation of *V. culicis* sporophorous vesicles in the mosquito gut. 4e. Both intense and lighter melanisation of spores. 4f. Uninfected mosquito gut, no sporophorous vesicles.

## Discussion

Being the first parasite to invade and colonise the mosquito host in the larval stage allows microsporidia to be at a distinct advantage over any later arrivals such as *Plasmodium* species. Not only has *V. culicis* been inside the mosquito host a good six to ten days before the arrival of any *P. berghei*, but it has also made the host’s environment its own and created a habitat to suit its own abundant growth and development. By the time the second invader arrives, *V. culicis* has not only depleted the host’s resources (Rivero A et al 2007), but changed the host’s internal environment, which is taken over by the unchecked multiplication of *V. culicis* spores. It should be considered surprising that anything at all could possibly grow and develop in mosquitoes under the conditions presented by *V. culicis* multiplication. That day six, *V. culicis* infected hosts could harbour double infections, albeit at low levels (average 3.3% of hosts found with oocysts) and not early, day two infected hosts, where spore production would have gained considerable hold, suggests that the intensity of *V. culicis* infection plays a significant part in the suppression of later *P. berghei* infection.

The results obtained here are similar to the very low levels of *Plasmodium* found in doubly infected hosts in the early studies on *Anopheles coluzzii* and *Nosema* undertaken by Fox RM and Weiser J (1959). They are slightly different to those obtained by Bargielowski I & Koella JC (2009) where the levels of *P. berghei* infection in two day old doubly infected hosts were higher (58.5%, compared to 0% from day 2 infected hosts from both experiments, here). This could be due to the different isolates of *V. culicis* used or the fact that fluorescent *P. berghei* which are clearly differentiated from microsporidian spores were unavailable for use in Bargielowski I & Koella JC’s (2009) study, or some other unknown combination of factors. Concordant to both studies, is that infecting early (day 2 post emergent) larvae, results in later *Plasmodium berghei* oocyst suppression in the adult stages of *Anopheles coluzzii*.

Direct comparisons cannot be drawn with other studies where double infections have been considered, for example in *Aedes aegypti* doubly infected with *V. culicis* together with the protozoan *Ascogregarina culicis* (Fellous S and Koella JC 2009); as there was no time lapse between infections, both parasites being larval infectors, were introduced to the host at the same time. Despite the difference, Fellous S and Koella JC’s 2009 study demonstrated the effect of increasing the infection level of *V. culicis* on the second parasite, *Ascogregarina culicis*, a result similar to that demonstrated here; that the higher levels of *V. culicis* offered by longer development times in day 2 infected larvae resulted in differential developmental success of the second malaria parasite. Here, there was by necessity a time lag between the two infections, microsporidians were introduced at two different larval ages, allowing for different developmental times and hence increases in parasite burdens of *V. culicis*; followed by malaria parasites in the adult stage.

Co-infection did not lead to a speeding up in the development of the *Plasmodium* parasite, as no sporozoites were seen to develop early. *P. berghei* sporozoites are known to egress from oocysts 11 to 12 days post infection (Klug & Frischknecht, 2017), none were observed. It would be expected that a longer period of observation should detect sporozoites from the controls, but the indication here was that co-infection did not hasten the development of *P. berghei* oocysts to sporozoites; as none were seen to develop early (before day 12 post infection) from doubly infected mosquitoes.

Margos G et al (1992) concluded that a lack of/competition for nutrition, was only partly responsible for the inhibition of *Plasmodium* in *Nosema* infected mosquitoes. However, transformation of the ookinete to oocysts is not completed in *Plasmodium* unless a range of nutrients are present (Carter V et al, 2007). In addition, transformation to the oocyst stage also required a change in pH, a fairly alkaline to neutral pH and in vitro, transformation was bicarbonate dependant (Carter V et al, 2007). *V. culicis* can germinate in a range of pH depending on the isolate, from a fairly acidic pH of 6.5, to 7 and 9.5 (Undeen AH 2007, Tsang KR et al 1982). Unless microsporidia affect the pH of the mosquito gut, pH should also not pose an issue to the development of either parasite, but nutrition might. Investigators have noted a reduction in some metabolites with microsporidian infection. For example, alkaline phosphatase which is known to promote gut health, nutrient absorption and prevention of bacterial invasion in mammalian guts, was shown to have reduced after *Nosema ceranae* infection in the honey bee *Apis mellifera*, indicating that infection by this microsporidian led to a reduction in gut protection and host health (Dussaubat C et al, 2012). Further studies on the nutritional requirements of *Plasmodium* in double infections are required as metabolite stores are severely limited in the ookinete and insufficient to complete transformation to the oocyst stage (Carter V et al, 2007). Carter’s study (2007) clearly indicated that transformation of the ookinete to oocysts required the correct pH, the presence of bicarbonate and the presence of a range of nutrients and all of these factors need to be considered in *V. culicis* infected hosts.

Fox RM and Weiser J (1959) suggested that the low/absence of *Plasmodium* oocysts in doubly infected hosts was due to the fact that *Nosema* infection had so disintegrated the mosquito gut that *Plasmodium* ookinetes could not possibly find suitable lodging in the tissue, a physical/mechanical issue that is seen from figures 3 & 4 from this study. Crossing of the gut wall has been described in other microsporidian species, for example, *Tubilinosema kingi* in *Drosophila melanogastor* (Vijendravarma RK 2008). Whilst the guts examined here were mostly intact (as seen under the light microscope), it is possible that in cases where the inner layers of cells lining the gut were damaged by *V. culicis*, allowed the sporophorous vesicles to protrude out and come into contact with the haemocele, resulting in a negative impact on both the parasites and the host. The extent of such damage can only be verified by electron microscope studies, to examine the inner gut cell linings at different infection levels.

The photographs (Figures 3 & 4) presented here show that *V. culicis* is able to cross the mosquito gut wall as evidenced by the sporophorous vesicles developing outwards from the gut wall into the haemocoel and in intimate contact with the cells lining the gut. However, this was only seen from mosquito guts heavily infected with microsporidia. It does seem that a substantial infection (as would be the case from early infected larvae) with microsporidia is necessary before it can have the desired effect of allowing no *Plasmodium* to develop. Although attempts have been made (Margos G et al 1992), it is difficult to quantify the exact or minimal level of microsporidian infection required that could halt plasmodium development.

As with microsporidia infecting an insect gut, malaria parasites have to pass through the peritrophic matrix (also known as the peritrophic membrane), which is a chitinous sack that facilitates digestion and protects the midgut epithelium lining the mosquito gut. It is known that malaria parasites have developed a specific mechanism, utilising a chitinase, to traverse the peritrophic matrix, before invasion of the midgut epithelium (Shahabuddin M et al 1993). Being intracellular parasites, microsporidians also invade the gut wall (Maddox JV et al 2000, Tanada Y & Kaya HA 1992, Becnel & Andreadis 1999). However, more needs to be known about how microsporidians get past the protective peritrophic matrix to get to the cells lining the gut wall.

Fox RM and Weiser J (1959) reported that the peritrophic matrix was not found in mosquitoes infected with Nosema. But it should be remembered that the formation of the peritrophic matrix is first initiated in the larval stage in *Anophelines* (Wigglesworth VB, 1930; Terra WR 2001) and parasites such as Nosema might influence peritrophic matrix formation. Microsporidian parasites would have had a much longer association with the peritrophic matrix (upwards of 12 days longer in early infected larvae and 8 days longer in later infected larvae) than malaria parasites. It is unlikely that microsporidia could use chitinase like *Plasmodium* to traverse the peritrophic matrix, as the microsporidian wall is made up of chitin and would therefore be destroyed by chitinase. The absence of the peritrophic matrix in microsporidian infected mosquitoes, suggests that microsporidia are either utilising it in some way, or preventing its formation by the secretary cells. It could also be the sheer pressure of intense parasitism by multiplying spores, that might influence peritrophic matrix formation and requires further examination.

Carter V et al (2007), have demonstrated that the cells of the basal lamina in the mosquito gut wall were not required to trigger ookinete to oocysts transformation for in vitro development of *Plasmodium* oocysts. Whilst the gut wall and hence basal lamina are not essential for oocyst development at least in vitro, it may be important in vivo, to protect oocysts from being melanised (Warburg A et al, 2007). The peritrophic membrane has been found to protect *Plasmodium* parasites by preventing mosquitoes from mounting an immune response to any invaders including *Plasmodium* (Kumar S et al, 2010). Hence, the absence of the peritrophic matrix due to microsporidian infection could well have a negative impact on *Plasmodium* species. If the peritrophic matrix protects *Plasmodium* species from the immune reaction of the host, it should in theory also protect microsporidian species and its absence should be catastrophic for both parasites, this too requires investigation. Melanisation of microsporidian spores was indeed seen here, Fig 4 c, d & e, however it was patches of melanisation and did not extend to all of the spores. The aim of this investigation was not to examine melanisation, but its presence was undeniable. It is evident that the gut wall was differentially damaged in some hosts allowing the haemocoel contents to gain entry into the gut wall and vice versa, to allow the gut contents, hence microsporidian spores to seep out, affording a route into the haemocoel leading to melanising of spores in some guts, Figure 4 c, d & e. However, contrary to previously held views, melanogenesis is not solely found in the haemocoel, where specialist immunogenic cells exist. Evidence points to sophisticated melanogenic activity within the gut; the cell wall of the gut does not need to be breached for melanisation to occur (Whitten MMA & Coates CJ, 2017). Further research is required on the reported absence of the peritrophic membrane in microsporidian infected mosquitoes as indicated by Fox RM and Weiser J (1959) and how, or if, this affects suppression of secondary infections and any parasite melanisation. This should lead to better understanding of the *Plasmodium* suppressive effects of microsporidia.

The melanisation of *V. culicis* spores as seen in the photographs (Figures 3 & 4) indicates that not all of the spores could be melanised, the response was not adequate to halt all spore production. Patches of both light and darker areas of melanisation was seen, as well as areas of unaffected spores (at least as seen by eye). The larval host was unable to sustain the melanisation attack or the parasite was able to evade it and only some of the spores, or some patches of spores were melanised (Figures 4d & 4e). More likely is that the rate of spore production is faster than the capacity of the numbers of haemocytes present or any other immunogenic gut process to melanise them. An initial dip in the numbers of spores produced was seen from older larval infections (Vyas-Patel N, 2020) indicating that older larvae may have a greater capacity to hinder spore production at the beginning of infections, compared to younger larvae. It would be advantageous to know if there was a difference in the numbers and types of haemocytes in younger compared to older infected hosts and to compare spore melanisation from younger and older infected larvae. Furthermore, to determine if the genes involved in the antimicrobial defence system which is known to regulate and defend mosquitoes against both bacterial and malarial parasites (Dong Y et al, 2006; Cirimotech CM 2010) were also similarly involved in the defence against *V. culicis*. Antimicrobial peptides are known to be a feature of microsporidian infections in a number of different hosts. Infection with microsporidians typically induced a strong host transcriptional response and up-regulation of a suite of different antimicrobial peptides (Jarkass THE & Reinke AW 2020). Dussaubat C et al (2012), demonstrated that the microsporidian *Nosema ceranae* silenced a significant number of genes involved in cell signalling, leading to an inhibition of renewal of intestinal cells in their honey bee hosts. Jarkass THE & Reinke AW (2020) and Dussaubat C et al (2012), stressed the importance of studying reactive oxygen species (ROS) which are efficient antimicrobial molecules and produced by insect guts as a general immune response to infection by microorganisms. All of the above requires examination in *V. culicis* infections of *Anophelines*.

Other avenues to explore would be to ascertain the biting capability of co-infected and singly infected mosquitoes. However, this would naturally give rise to ethical concerns. Hulls RH, (1971) reported that *P. berghei* sporozoites from *Nosema* infected mosquitoes were less infective compared to controls when inoculated into mice, but this does not address the natural biting capability of doubly infected mosquitoes. Until then, any control strategy aimed at using microsporidia with a view to halting the development of *Plasmodium* development should therefore target early, newly hatched larvae.

Efforts aimed at reducing malaria parasites in the adult mosquito clearly requires a critical level of microsporidian infection to be present within the host, at least in the case examined here. Microsporidian infections reduced the size of individual malaria parasites (Bano L 1958, Bray RS 1958), but not enough is known about the capacity and transmission capability of smaller sized malaria parasites to cause disease. The parasites *V. culicis* and *P. berghei* were used in this study because cultures of both were established in the laboratory and could readily and safely be used. *V. culicis* is horizontally transmitted; maintaining very high levels of the parasite in the field may require regular, inundative, releases. Vertically transmitted microsporidia might negate the requirement of regular releases of microsporidia but the level of microsporidian infection required to attain malaria parasite suppression would need to be assessed. High levels of malaria parasite suppression was seen in the naturally occurring, vertically transmitted microsporidian, *Microsporidia MB*, from *Anopheles arabiensis* infected mosquitoes (Herren JK et al 2020). This suggests that if the spores are vertically transmitted, the levels of microsporidian infection required to interfere with oocyst development may not matter and this could easily be assessed. Generalizations cannot be made however, as in the horizontally transmitted *Nematocida parisii* in the nematode host *Caenorhabditis elegans*, it was found that infection resulted in host offspring with enhanced immunity to the microsporidian. In this case, although not a mosquito host, the immune response inherited from the parent, only lasted for one generation and was dose dependant (Willis AR et al, 2021). Each individual parasite species and host system needs to be individually studied and assessed.

An interaction exists between the two parasites with the sheer numbers of microsporidia exerting a strong negative impact on the numbers of plasmodium that can develop in doubly infected hosts. This is evidenced by the presence of oocysts from a few mosquitoes infected with *V. culicis* later on (day 6 post hatching) compared to early infected mosquitoes (day 2 post hatching) where *P. berghei* development was completely halted. The numbers of *V. culicis* spores, hence the intensity of microsporidian infection, plays a major part in the suppression of *P. berghei* development in doubly infected mosquitoes.

## Acknowledgments

This project was funded by the Daphne Jackson Trust and Imperial College; both are gratefully acknowledged. Special thanks are due to Prof. Elizabeth U Canning and Prof Jacob Koella for discussion of the photographs and the study.

